# Sequence-Specific Targeting of GC-Rich Gene Loci by Parallel Triplex-Forming Oligonucleotides Containing a Modified Nucleobase

**DOI:** 10.64898/2026.07.30.741700

**Authors:** Ruolin Ma, Michael Brazzill, Dylan Justice, Nicholas Buckham, Cen Chen, Shuichi Hoshika, Steven A. Benner, David A. Rusling

## Abstract

Targeting GC-rich gene loci is a major challenge owing to their high duplex stability, repetitive sequence composition, and propensity to adopt alternative DNA structures. Triplex-forming oligonucleotides (TFOs) provide a programmable strategy towards the recognition of GC-rich DNA, but their application is restricted by the limited recognition capabilities of natural nucleobases in a cellular setting. Here, we overcome this barrier using parallel-binding TFOs containing the synthetic nucleobase 6-amino-5-nitropyridin-2-one (Z), which enables pH-independent recognition of G-C base pairs. Using two structurally distinct regulatory elements within the *MYC* promoter, we show that Z-modified TFOs form stable, sequence-selective triplexes that repress promoter activity by 50-80% in both episomal reporter assays and at endogenous gene loci. Notably, the greatest repression was observed at a GC-rich quadruplex-forming element that functions as a structural hub for transcription factor recruitment. To our knowledge, this represents the first demonstration that a simple nucleobase modification alone is sufficient to enable parallel-binding TFOs to repress expression of an endogenous gene, establishing a general strategy for targeting GC-rich regulatory elements through programmable DNA recognition.

**TOC graphic:** 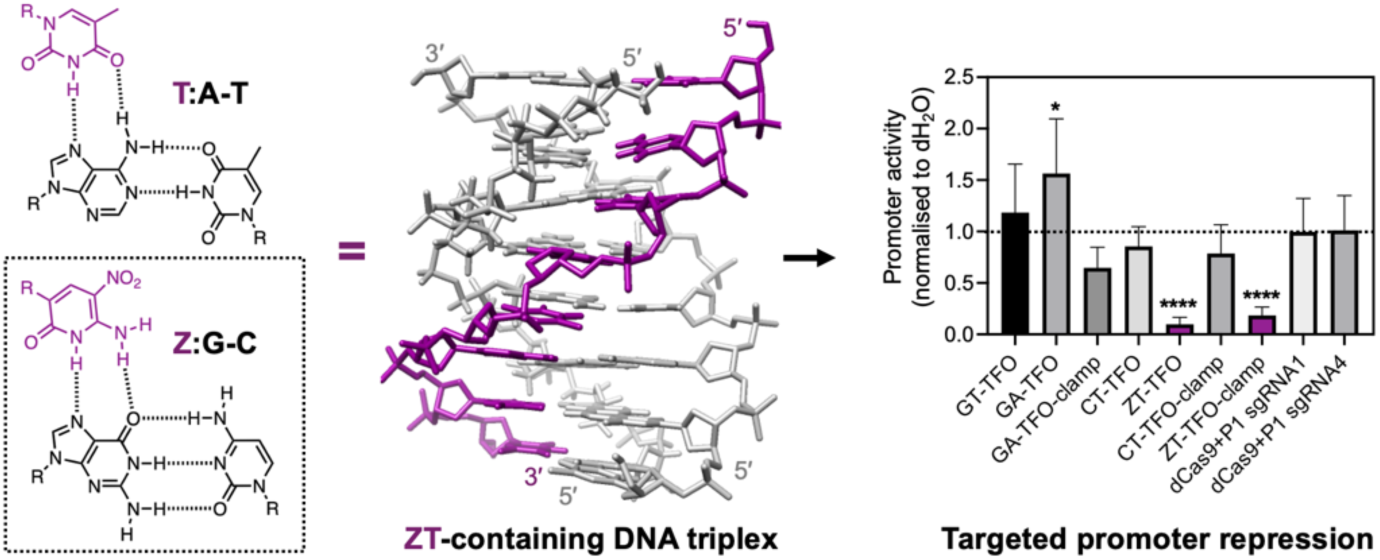

## INTRODUCTION

GC-rich DNA represents an important yet persistently challenging target for sequence-selective ligands, oligonucleotides, and programmable DNA-binding proteins. These sequences are enriched within promoters, CpG islands, and other regulatory elements, where they contribute to transcriptional and epigenetic control. However, targeting these loci is hindered by genomic context and intrinsic physicochemical properties, including high duplex stability, repetitive sequence composition, and structural polymorphism. For example, the elevated thermodynamic stability of GC-rich duplexes can impede targeting strategies that rely on strand separation;[1, 2] repetitive tracts reduce selectivity by generating multiple, closely related binding sites;[3] and the propensity of these sequences to adopt alternative secondary structures, such as G-quadruplexes and i-motifs, can further occlude target binding.[4, 5] Consequently, GC-rich gene loci remain critically important biological targets that are intrinsically difficult to engage with high efficiency and selectivity.

Triplex-forming oligonucleotides (TFOs) provide a programmable strategy towards targeting GC-rich DNA because they recognize oligopurine-oligopyrimidine tracts directly within the major groove without requiring strand separation and/or protein intermediaries (**Figure 1**).[6, 7] Alternatively, TFO-clamps can be designed to bind single-stranded DNA regions by forming both Hoogsteen and Watson-Crick (W-C) hydrogen bonds, generating stable triplex complexes.[8–12] Because TFOs target extended sequences, they offer significantly greater sequence selectivity than small-molecule ligands and have shown potential as transient gene-targeting tools, modulating gene expression *in vitro*,[13–15] in cell culture, and in animal models.[16, 17] Moreover, because triplexes can exhibit comparable or superior stability to alternative DNA structures such as G-quadruplexes, TFOs and TFO-clamps offer a means to dynamically compete with their formation.[18, 19]

**Figure 1.**
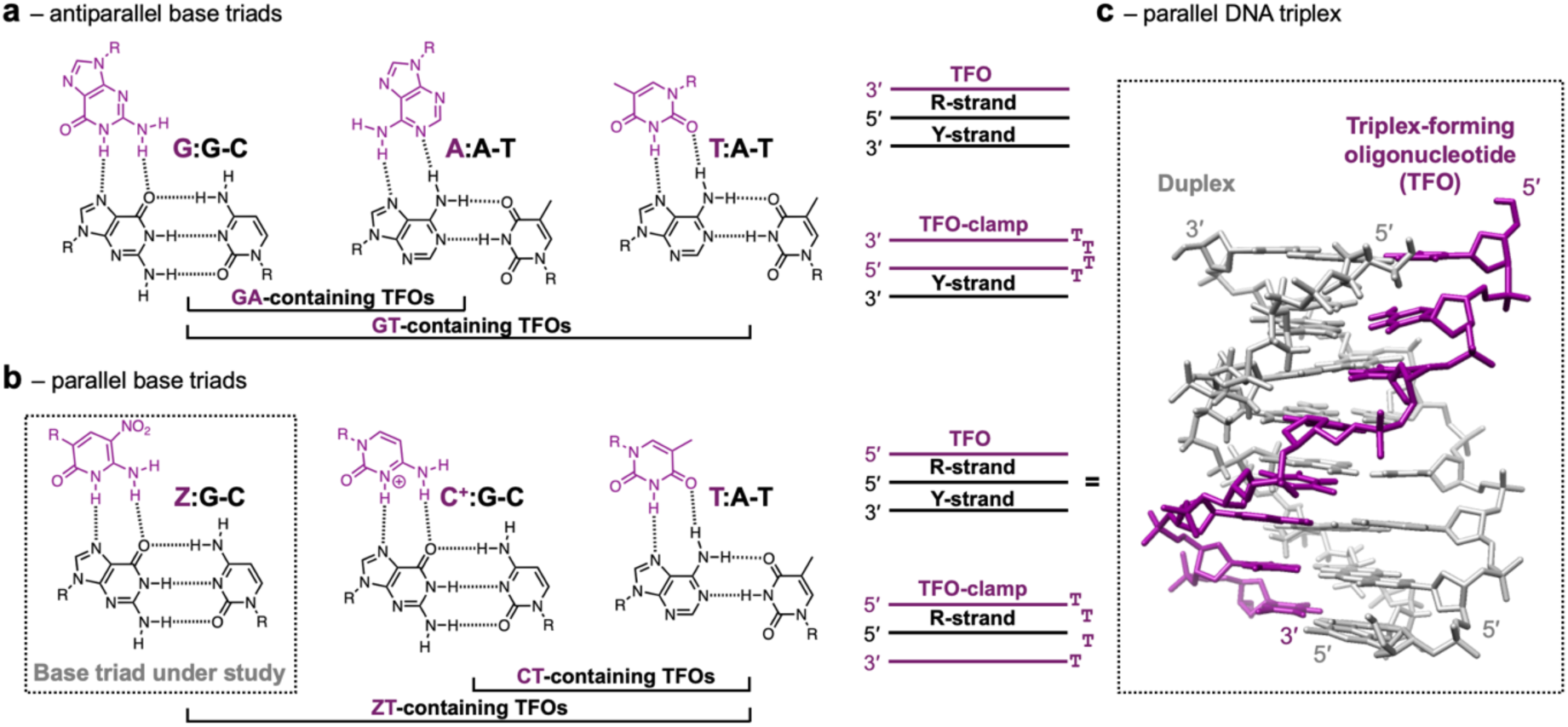
Triplex formation by canonical and modified base triads. **(a-b)** Chemical structures and strand orientations of TFOs and TFO-clamps forming antiparallel and parallel base triads. Here, X:R-Y is used to denote a triad, where X is the third strand base and R-Y is the target duplex base pair. Base pairs are shown in black, while the TFO and TFO-clamps are shown in purple. TFOs target intact duplexes, while antiparallel or parallel TFO-clamps bind single-stranded regions of DNA by forming both Hoogsteen and W-C interactions upon strand separation. The pH-independent parallel Z:G-C triad investigated in this study is highlighted by the dashed box. **(c)** NMR structure of a parallel DNA triplex formed by C^+^:G-C and T:A-T triads, with the W-C duplex shown in grey and TFO in purple (PDB code: 1D3X).

However, the triplex-forming capacity of oligonucleotides composed of natural nucleobases is constrained by several sequence and structural requirements that limit their efficacy in a cellular setting. GT- or GA-containing TFOs typically bind antiparallel to the purine strand of the target duplex through formation of G:G-C and either T:A-T or A:A-T base triads (**Figure 1a**).[20, 21] However, the resultant triads are not isostructural, leading to backbone distortions between adjacent triads that reduce triplex stability,[22] and can also stimulate DNA repair and recombination.[23] Moreover, purine-rich oligonucleotides readily self-associate into competing secondary structures, including G-quadruplex and related assemblies, decreasing the effective concentration of the active species and complicating interpretation of biological activity.[24–26] Such folded structures can also be intrinsically toxic to living cells.[26, 27] By contrast, CT-containing oligonucleotides bind in a parallel orientation *via* C^+^:G-C and T:A-T base triads (**Figure 1b** and **1c**).[28, 29] Since the constituent base triads are isostructural they are inherently more stable than their antiparallel counterparts and also less prone to unwanted mutagenesis;[23] however, their formation depends heavily on the protonation of cytosine at N3 (p*K*_a_ ≈ 4.5), restricting efficient binding to mildly acidic conditions (< pH 6.0).

Consequently, considerable effort has been devoted to the development of base-modified TFOs that preserve the favorable geometry of parallel triplex formation while removing the pH dependence of cytosine-containing third strands.[7, 30] Among the most promising is 6-amino-5-nitropyridin-2-one,[31, 32] an uncharged *C*-glycoside mimic of protonated cytosine that enables stable recognition of G-C base pairs at physiological pH when used alongside T:A-T base triads (**Figure 1b**).[33] Importantly, the presence of a proton at N3 reduces the propensity of Z to form W-C base pairs with guanine in single-stranded nucleic acids, suggesting Z-containing TFOs might exhibit improved triplex selectivity.[33] Although the Z nucleobase has shown exceptional triplex-forming capacity *in vitro*,[34] and alongside other nucleobases can be used to target duplex sequences containing non-standard base pairs,[35] its application to gene-targeting and transcriptional regulation has not yet been explored.

Here, we therefore evaluate the anti-gene potential of parallel-binding ZT-containing TFOs directed against two structurally distinct GC-rich regulatory elements within the *MYC* promoter. Importantly, we benchmark their performance against unmodified GT-, GA-, and CT-containing TFOs and TFO-clamps, as well as catalytically inactive dCas9, under identical experimental conditions. By targeting oligonucleotide duplexes, supercoiled plasmids, and endogenous DNA, we show that ZT-modified parallel-binding TFOs form stable, sequence-selective triplexes with GC-rich DNA and repress promoter activity and endogenous *MYC* expression. To our knowledge, this represents the first demonstration that a simple nucleobase modification alone is sufficient to enable parallel-binding TFOs to repress expression from an endogenous human gene locus, providing a straightforward strategy for programmable recognition and regulation of GC-rich genomic DNA.

## MATERIALS & METHODS

### Reagents

Unmodified oligonucleotides were purchased from Sigma-Aldrich (Dorset, UK), and Z-containing oligonucleotides were synthesised by Firebird Biomolecular Sciences (Alachua, FL, USA) as previously described.[37] EnGen® sgRNA Synthesis Kit, Monarch® RNA Cleanup Kit, SpyCas9 nuclease, EnGen® SpRY Cas9 and NEBuffer r3.1 were obtained from New England Biolabs (Ipswich, MA, USA). Lipofectamine™ 3000, Oligofectamine™ and PrestoBlue™ Cell Viability Reagent were obtained from Invitrogen (Thermo Fisher Scientific, Paisley, UK). Opti-MEM, Dulbecco’s Modified Eagle Medium (DMEM), fetal bovine serum (FBS), and penicillin-streptomycin were obtained from Gibco (Thermo Fisher Scientific, Paisley, UK). GelRed was obtained from Biotium (Freemon, CA, USA). Luciferase Assay System was purchased from Promega (Madison, WI, USA). RNeasy Mini Kit was obtained from Qiagen (Hilden, Germany), High-Capacity cDNA Reverse Transcription Kit from Applied Biosystems (Thermo Fisher Scientific, Paisley, UK), and SsoAdvanced Universal SYBR Green Supermix from Bio-Rad (Hercules, CA, USA). Primary antibodies used were anti-c-MYC (Y69; Abcam, Cambridge, UK) and anti-GAPDH (6C5, Abcam, Cambridge, UK), together with IRDye® 680RD goat anti-rabbit IgG and IRDye® 800CW goat anti-mouse IgG secondary antibodies (LI-COR Biosciences, Lincoln, NE, USA). Circular dichroism spectra were acquired using a Chirascan VX spectropolarimeter (Applied Photophysics, UK), UV melting experiments using a Shimadzu UV-3600 spectrophotometer (Shimadzu, Japan), and fluorescence and luminescence measurements using a SpectraMax i3x plate reader (Molecular Devices, San Jose, CA, USA).

### Biological Resources

HEK293 and HCT116 cell lines were obtained from the University of Portsmouth Cell Bank (Portsmouth, UK) and maintained under standard culture conditions. The *MYC* promoter reporter plasmid (Del4) was a gift from Bert Vogelstein (Addgene plasmid #16604).[38]

### Websites/Databases

The TFO Target Sequence Search tool (http://utw10685.utweb.utexas.edu/tfo/)[36] was used to inform oligonucleotide design, with target sequence composition and length chosen to ensure that each target was predicted to occur only once within the human genome. Oligonucleotide sequences used in this study are shown in **Table S1**.

### Electrophoretic mobility shift assay (EMSA)

Interactions of the TFOs with P1 and P2 duplexes or their constituent strands were determined by native EMSA at different pH. Oligonucleotides were prepared in 10 mM sodium cacodylate containing 10 mM magnesium chloride at either pH 5.0 or pH 7.0. The final duplex concentration was 1 μM and the TFO concentration varied between 10 μM and 0.01 μM in a total volume of 20 µL as indicated. Duplexes were annealed by heating to 95°C for 5 min followed by slow cooling to room temperature over 2 h, after which TFOs were added and the samples equilibrated for >16 h at 4°C. Samples were then subjected to non-denaturing polyacrylamide gel electrophoresis at 100 V for ∼2 ½ h in 40 mM tris-acetate running buffer at either pH 5.0 or pH 7.0 as indicated. Gels were post-stained with GelRed and visualised using a Gel Doc XR+ Imaging System.

### Circular dichroism (CD)

CD spectra for the duplexes and triplexes were determined using a Chirascan VX spectrophotometer. Oligonucleotides were prepared in 10 mM sodium cacodylate containing 10 mM magnesium chloride at either pH 7.0 or pH 5.0. The final duplex concentration was 5 μM and final TFO concentration was 10 μM in a total volume of 300 µL. Duplexes were annealed by heating to 95°C for 5 min followed by slow cooling to room temperature over 2 h, after which TFOs were added and the samples equilibrated for >16 h at 4°C. Spectra were collected between 320–200 nm, at 100 nm/min, 1 s response time, 1 nm bandwidth in Hellma synthetic quartz cuvettes with a 1 mm pathlength. Each spectrum was accumulated five times, averaged, and smoothed.

### Ultraviolet (UV) melting

Thermal melting profiles for the duplexes and triplexes were determined using a Shimadzu UV-3600 UV-Vis-NIR spectrophotometer. Oligonucleotides were prepared in 10 mM sodium cacodylate containing 10 mM magnesium chloride at pH 5.0 or pH 7.0. The final duplex concentration was 5 μM and final TFO concentration was 10 μM in a total volume of 100 µL. Duplexes were annealed by heating to 95°C for 5 min followed by slow cooling to room temperature over 2 h, after which TFOs were added and the samples equilibrated for >16 h at 4°C. UV melting profiles were recorded at 260 nm, 1 s response time, in a Shimadzu 8 series micro multi-cell with a 10 mm pathlength between 30°C and 90°C at a ramp rate of 0.2°C/min. Melting temperatures (*T*_m_) were determined from the first derivatives of each profile using the analysis software provided by the machine, and typically varied by less than 1°C between experiments. Mean values ± SEM were calculated from two independent measurements on distinct samples.

### sgRNA design and Cas9 assays

Guide RNAs (sgRNA) were generated using the EnGen® sgRNA Synthesis Kit according to the manufacturer’s protocol. In vitro transcription reactions containing 1 μM DNA template in a total volume of 20 μl were carried out at 37°C for 30 min. Reactions were immediately cooled on ice and brought to a final volume of 50 μl with nuclease-free water. DNA template was removed by addition of 2 μl DNase I followed by incubation at 37°C for 15 min. sgRNA were purified using the Monarch® RNA Cleanup Kit in accordance with the manufacturer’s instructions, and RNA concentration was determined by NanoDrop 2000 spectrophotometry.

For the Cas9 protection assay, 30 nM of the *MYC* promoter reporter plasmid (Del4) was incubated with TFO at final concentrations ranging from 10 to 0.1 μM in 1X NEbuffer r3.1 in a total volume of 3 μl. Samples were equilibrated for >16 h at 4°C. Cas9 nuclease from *S. pyogenes* or EnGen® SpRY Cas9 were assembled with ∼30 nM sgRNA according to the manufacturer’s instructions in a total reaction volume of 27 μl and incubated at room temperature for 10 min. The Cas9/sgRNA mixture was then added to each pre-equilibrated plasmid/TFO sample and incubated at 37°C for 1 h. Reactions were terminated by heating at 70°C for 10 min to inactivate Cas9. Samples were resolved on 0.8% agarose gels containing GelRed in standard 40 mM tris-acetate EDTA running buffer at 100 V for ∼1 ½ h. Gels were post-stained with GelRed and visualised using a Gel Doc XR+ Imaging System.

### Cell culture

HEK293 and HCT116 cells were cultured in Dulbecco’s Modified Eagle Medium (DMEM) growth media supplemented with 10% fetal bovine serum (FBS) and 1% penicillin-streptomycin. Cells were maintained at 37°C in a humidified incubator with 5% CO_2_ and used at passages <30.

### Luciferase activity assay

HEK293 cells were maintained in DMEM supplemented with 10% FBS at 37°C in a humidified atmosphere containing 5% CO_2_. Cells were seeded in duplicate in 96-well plate at a density of 2 x 10^4^ cells per well and allowed to adhere overnight. For transfection, 100 ng (*ca.* 0.5 nM) of the *MYC* promoter reporter plasmid (Del4) was pre-incubated with TFO at final concentration of either 67, 6.7 or 0.67 nM, or with vehicle dH_2_O control (RNase-free water) for >16 h at 4°C. The plasmid/TFO mix was then combined with Lipofectamine 3000^TM^ at a 1:1 ratio and incubated for 20 min at room temperature prior to addition to the cells. After 24 hours, luciferase activities were measured using a Luciferase Assay Kit and a SpectraMax i3x Microplate Reader. Luciferase activity was expressed relative to the untreated dH_2_O control. Experiments were performed with a minimum of three biological replicates. Statistical significance was determined by one-way ANOVAs followed by Tukey’s post hoc test.

### Quantitative real-time PCR (qRT-PCR)

HCT116 cells were seeded in 6-well plates at 2 x 10^5^ cells per well in DMEM supplemented with 10% FBS and allowed to adhere overnight. Before transfection, the culture medium was replaced with 800 μl DMEM. TFOs were diluted in Opti-MEM to final concentrations of 200, 120 or 20 nM, complexed with Oligofectamine^TM^ Reagent in a total volume of 200 μl, and added to the cells. After 6 hours, 500 μl DMEM supplemented with 30% FBS was added to each well.

At 24 hours post-transfection, cells were washed with 1X PBS and lysed, and total RNA was extracted using a Qiagen RNeasy Mini kit according to the manufacturer’s instructions. Total RNA (1000 ng) was reverse transcribed into cDNA using a High-Capacity cDNA Reverse Transcription Kit according to the manufacturer’s instructions. cDNA synthesis was performed on a Bio-Rad T100 Thermal Cycler under the following conditions: 25°C for 10 min, 37°C for 120 min, 85°C for 5 min, followed by a 4°C hold. Quantitative PCR was carried out using SsoAdvanced Universal SYBR Green supermix with primers designed specific for *MYC*, *GAPDH* and *β-actin* mRNA (**Table S1**). The qPCR reactions were thermocycled on a Roche LightCycler 96 instrument (Roche) using following protocol: pre-incubation, 95°C for 60 s (1 cycle), followed by 2 step amplification, 95°C for 15 s and 60°C for 60 s (45 cycles), finally melting step, 60°C for 15 s and 95°C for 1 s. Relative expression levels were calculated using the 2^−ΔΔCt^ method, normalised to the mean of *GAPDH* and *β-actin*, and expressed relative to the vehicle control (dH₂O). Fold change values were plotted as log₂-transformed data. Experiments were performed with a minimum of three biological replicates. Statistical significance was determined by one-way ANOVAs followed by Tukey’s post hoc test.

### Western blot (WB)

HCT116 cells were cultured and transfected as described above for RT-qPCR, except that cells were treated with oligonucleotides at either 2,500, 250, 25 nM or vehicle control (dH₂O) only. After 48 hours post-transfection, cells were washed twice with 1X PBS and harvested using 1X RIPA lysis buffer supplemented with 1 X Halt^TM^ protease and phosphatase inhibitor cocktail, EDTA-free. Cell debris was removed by centrifugation for 10 min at 13,300xg and 4°C. The total protein concentration of the cleared lysate was then measured by Pierce BCA Protein assay kits at 560 nm using a SpectraMax i3x Microplate Reader. A total of 25 µg protein was then loaded on a 4% / 10% polyacrylamide SDS-PAGE gel and transferred to an Amersham Protran 0.45 μm NC nitrocellulose blotting membrane for 30 min and 300mA using a semi-dry Trans-Blot Turbo Transfer System. The membrane was blocked (5% ski-milk in TBS-T) and incubated with primary antibodies (1: 20,000 Anti-GAPDH antibody – 6C5, and 1:1000 Anti-c-MYC antibody - Y69, Abcam) overnight at 4 °C. The membrane was then washed in 1X TBS-T three times at room temperature and then incubated with secondary antibodies (1: 10,000 IRDYE 680RD Goat Anti-Rabbit IgG and 1: 10,000 IRDYE 800CW Goat Anti-mouse IgG) for 1 h at room temperature. The membrane then washed by 1X TBS-T three times. The signal was developed using LI-COR Odyssey CLx imager. Quantification was performed using the ImageJ software. Relative MYC protein band intensity was normalized to GAPDH and then normalized to dH_2_O control. Experiments were performed with a minimum of three biological replicates. Statistical significance was determined by one-way ANOVAs followed by Tukey’s post hoc test.

### Cell viability assays

Cell viability was assessed using PrestoBlue™ Cell Viability Reagent according to the manufacturer’s instructions. Cells were seeded in duplicate in 96-well plates at 2 x 10^4^ cells per well in 100 μl DMEM supplemented with 10% FBS and allowed to adhere overnight prior to transfection. Immediately before transfection, the medium was replaced with 70 μl DMEM supplemented with 1% FBS. At 48 h post-transfection, 8 μl PrestoBlue™ reagent was added directly to each well and incubated for 30 min at 37°C. Fluorescence was then measured using a SpectraMax i3x microplate reader with excitation at 560 nm and emission at 590 nm. Experiments were performed with a minimum of three biological replicates and viability values were normalised to the vehicle control (dH₂O). Statistical significance was determined by one-way ANOVA followed by Tukey’s post hoc test.

## RESULTS

### Oligonucleotide design and benchmarking

The *MYC* gene is a widely used model for benchmarking oligonucleotide-mediated gene-targeting in both cell-free and cellular systems.[19, 39–47] Its promoter and transcribed regions contain multiple oligopurine-oligopyrimidine tracts supporting triplex formation that could sterically block transcription initiation or elongation. The best-characterised lie upstream of the P1 and P2 promoter transcription start sites and correspond to sequences 27- and 23-base pairs in length, respectively (**Figure 2**). These regulatory elements participate in a complex network of interactions with nuclear proteins that modulate *MYC* transcription.[48] Notably, the P1 sequence also exists in a dynamic equilibrium between duplex, triplex, and quadruplex conformations,[49, 50] with the latter implicated as a structural hub for transcription factor recruitment.[51]

**Figure 2.**
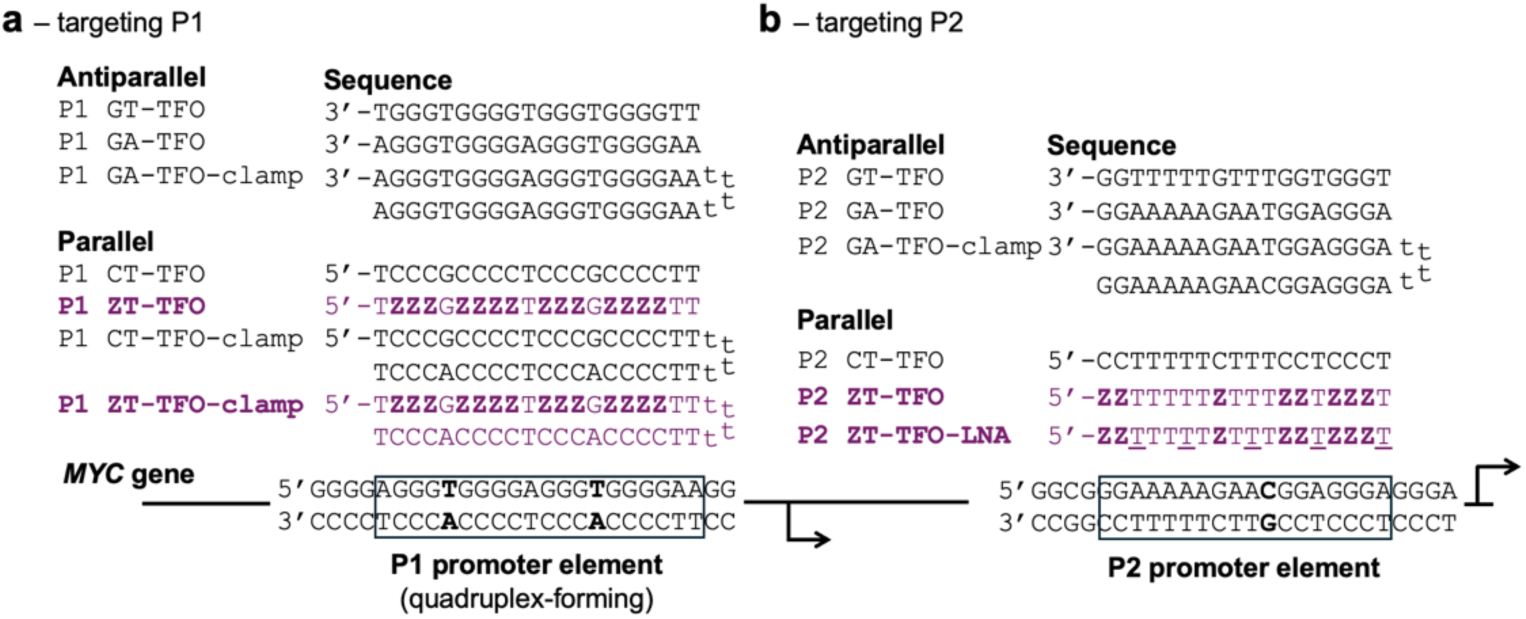
Targeting P1 and P2 promoter elements in the *MYC* promoter by triplex formation. **(a-b)** Sequences of ZT-containing TFOs and TFO-clamps targeting oligopurine-oligopyrimidine tracts in the P1 and P2 promoter elements and benchmarked against equivalent parallel CT-containing and antiparallel GT- and GA-containing oligonucleotides. Target duplex sequences and base pairs are shown in black, the TFO target sequence boxed, and pyrimidine-purine inversion sites are highlighted in bold. The ZT-containing TFOs and their constituent bases are shown in purple, with the positions of the Z nucleobase indicated in bold. Underlined Ts in the P2 ZT-TFO-LNA oligonucleotide represent LNA-modified residues.

To date, the P2 promoter element has proven more amenable to triplex targeting than P1, owing to its lower GC-content and likely duplex-only conformation.[19, 39–41, 43–45] Previous studies targeted the complete 23-bp oligopurine tract using antiparallel purine-rich GT- or GA-containing TFOs. Here, we instead selected a truncated 18-bp region predicated to occur just once in the genome, allowing shorter oligonucleotides to be used without compromising sequence selectivity (**Figure 2b, boxed)**. Two parallel TFOs containing the Z nucleobase were prepared: one comprising Z and T alone (P2 ZT-TFO) and a second incorporating five dispersed LNA-thymidine residues to establish whether enhancements to TFO affinity correlated with improved bioactivity (P2 ZT-TFO-LNA) (**Figure 2b, Table S1**).[52] These were benchmarked against four unmodified control oligonucleotides: an equivalent parallel CT-containing TFO (P2 CT-TFO), antiparallel GT- and GA-containing TFOs (P2 GT-TFO and GA-TFO), together with a GA-TFO-clamp designed to engage the pyrimidine (Y) strand through combined Hoogsteen and W-C interactions (P2 GA-TFO-clamp) (**Figure 2b, Table S1**). The single C-G interruption within the target sequence was accommodated using thymine to generate a T:C-G triad upon triplex formation.[53]

The P1 promoter element represents a more challenging target because its 27-bp sequence contains five G-tracts associated with secondary structure formation.[49, 54] To challenge triplex formation under these conditions, we selected a central 20-bp region encompassing four of the five G-tracts (**Figure 2a, Table S1**). Two parallel ZT-containing oligonucleotides were prepared: one compromising of Z and T alone (P1 ZT-TFO) and a second clamp designed to form additional W-C interactions with the purine (R) strand to establish whether this conferred greater bioactivity (P1 ZT-TFO-clamp). These were benchmarked against five unmodified control oligonucleotides: a corresponding CT-containing TFO (P1 CT-TFO) and TFO-clamp (P1 CT-TFO-clamp), together with antiparallel GT- and GA-containing TFOs (P1 GT-TFO and P1 GA-TFO), and a final GA-TFO-clamp (P1 GA-TFO-clamp) (**Figure 2a, Table S1**). The two T-A interruptions within the target sequence were accommodated using thymine in antiparallel TFOs or guanine in parallel TFOs, generating T:T-A or G:T-A base triads, respectively.[55–57] For both P1 and P2, a highly modified Z-containing oligonucleotide served as a control to confirm that any observed effects were dependent on sequence-specific target recognition rather than the presence of the synthetic nucleobase itself (ZT-TFO-scrambled) (**Table S1**).

### Stable and selective triplex formation by ZT-containing TFOs targeting P1 and P2

We first investigated triplex formation between the oligonucleotides and synthetic P1 and P2 target duplexes by an electrophoretic mobility shift assay (EMSA) (**Figure 3a, 3b**). Importantly, binding was also assessed with the constituent purine (R) or pyrimidine (Y) strands of each duplex in isolation. This allowed us to distinguish *bona fide* triplex formation from triplex-independent interactions in later experiments. This comparison was particularly important for the repetitive P1 sequence, where unmodified purine- or pyrimidine-rich TFOs were expected to exhibit competing interactions with isolated duplex strands (*e.g.,* **Figure S1a**). In contrast, ZT-containing TFOs were not expected to form these unwanted interactions due to reduced stability of the G-Z base pair at neutral pH (*e.g.,* **Figure S1b**).[33]

**Figure 3.**
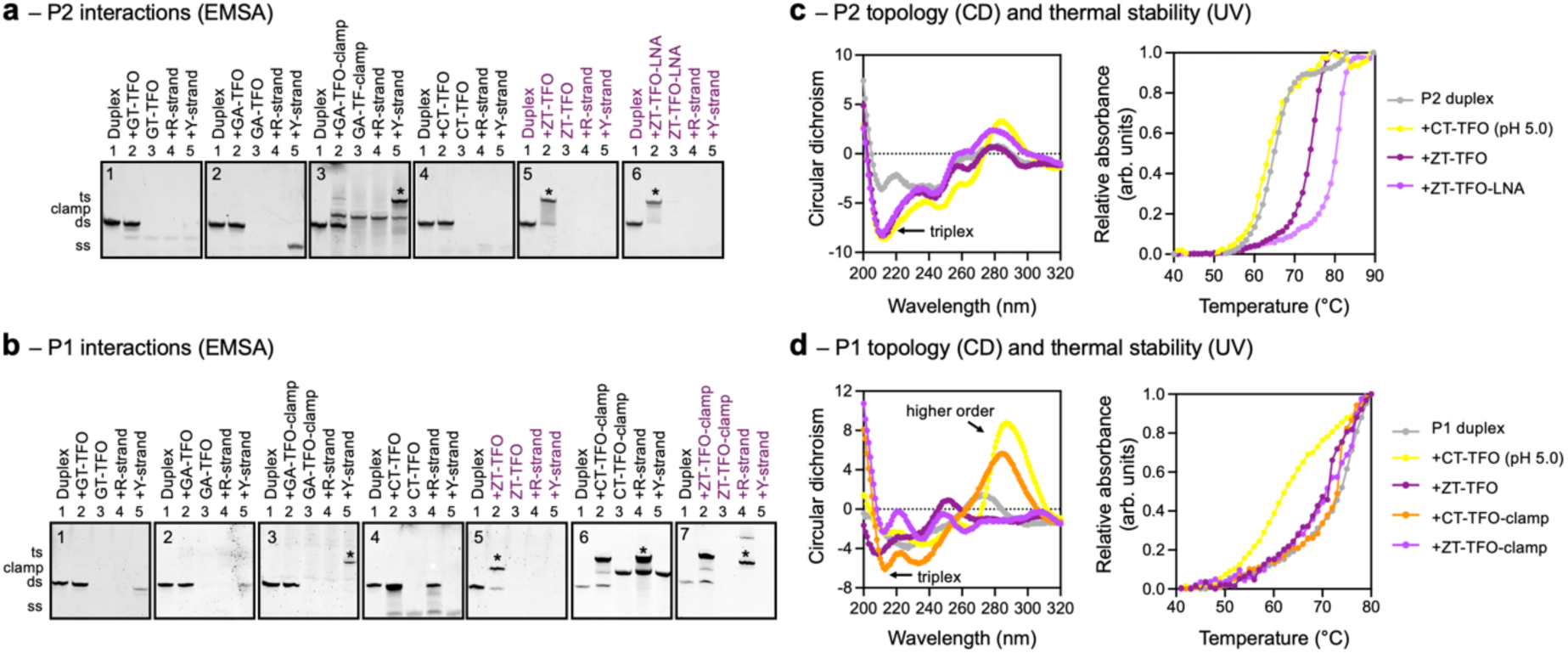
Stable and selective triplex formation by ZT-containing TFOs targeting P1 and P2. **(a-b)** EMSA analysis of oligonucleotide interactions. Oligonucleotides were annealed at pH 7.0 in a sodium cacodylate buffer containing 10 mM magnesium at a final duplex and TFO concentration of 1 μM and 2 μM, respectively. Complexes were resolved on a 20% non-denaturing polyacrylamide gel in standard tris-acetate running buffer and visualized by GelRed staining. Successful triplex formation is indicated by a mobility shift relative to the duplex-only control (asterisk), whereas unintended interactions between the oligonucleotides and constituent duplex strands (P1 only) migrate similarly to the duplex-only control. **(c-d)** CD and UV melting analysis of triplex samples. Oligonucleotides were annealed in a sodium cacodylate buffer containing 10 mM magnesium at final duplex and TFO concentration of 5 μM and 10 μM, respectively. CD spectra were collected between 320-200 nm and each spectrum represent an average of five accumulations. Triplex formation was evidenced by an increased negative peak at 210 nm compared to the duplex-only control (black arrow). UV melting profiles were recorded by measuring absorbance at 260 nm as the complexes were heated at a rate of 0.2 °C/min (n >2). Triplex formation is evidenced by either a shift in melting curve relative to the duplex-only control or the melting of the entire complex in a single process (P1 only).

We first evaluated targeting P2 which lacks repetitive GC-rich sequence elements (**Figure 3a**). Analysis of the gels revealed that none of the unmodified antiparallel GT- or GA-containing TFOs, nor the parallel CT-containing TFO, formed stable interactions with either the duplex (lane 1 vs 2; gel 1, 2, and 4) or individual duplex strands under the experimental conditions tested (lane 3 vs 4 or 5; gel 1, 2, and 4). The GA-containing TFO-clamp, however, produced a shifted species with the duplex, but substantially stronger binding to the isolated pyrimidine strand indicating that W-C interactions rather than stable triplex formation predominated (lane 5; gel 3). The CT-containing TFO failed to form a stable complex at concentrations up to 10 μM at pH 7.0 and exhibited detectable binding only by reducing to pH 5.0 (**Figure S2a**).

By contrast, both the ZT-containing TFO and its LNA-modified derivative formed discrete complexes at neutral pH (lane 1 vs 2; gel 5 and 6) and did not show any interaction with the isolated duplex strands alone (lane 3 vs 4 or 5, gel 5 and 6). Triplex formation by the ZT-containing TFOs was concentration-dependent as expected for an intermolecular interaction (**Figure S3a**).

We next evaluated targeting P1 which contains multiple repeated GC-tracts within its sequence (**Figure 3b**). Under the buffer conditions selected, and in the absence of supercoiling, duplex oligonucleotides were not expected to form competing structures. Indeed, samples produced a single species on the gel attributed to the formation of the intended duplex structure (*e.g.,* lane 1 of each gel). Analysis of samples in the presence of TFOs revealed that again none of the unmodified antiparallel GT- or GA-containing oligonucleotides, nor the parallel CT-containing oligonucleotide, formed detectable interactions with the pre-assembled duplex under these conditions (lane 1 vs 2; gel 1, 2, and 4). Instead, the GT- and GA-containing TFOs interacted with the pyrimidine strand alone, whereas the CT-containing TFO interacted with the individual purine strand alone, presumably through unwanted W-C base pairings (lane 4 or 5; gel 1, 2, and 4)(**Figure S1a**). As observed for P2, the CT-containing TFO formed detectable complexes only at acidic pH, but additional slowly migrating species also appeared, suggesting competing higher-order structures (**Figure S2b**). By contrast, the ZT-containing TFO formed a stable triplex at neutral pH and, due to the reduced capacity of Z to bind to the natural nucleobases, showed no interaction with the individual duplex strands alone (lane 1 vs 2-5; gel 5) (**Figure 3b, S3b**).

Similar experiments were performed with the corresponding CT- and ZT-containing oligonucleotide clamp derivatives, which were designed to form additional W-C interactions with the purine strand of the duplex (**Figure 3b**). This time, both the unmodified and modified oligonucleotides formed shifted species on the gels, likely reflecting the formation of triplex or partial triplex structures (lane 2 or 5; gel 6 and 7).

For both P1 and P2 targets, additional experiments revealed that the shifted species on each gel correspond to similar oligonucleotide architectures, as running the complexes side-by-side gave similar mobilities (**Figure S4**). To verify that the shifted species corresponded to triplex structures, the complexes were analyzed by circular dichroism (CD) (**Figure 3c, 3d**). All stable complexes formed by the ZT-containing TFOs, together with the CT-containing TFOs at pH 5.0 and both TFO-clamps, exhibited an enhanced negative band near 210 nm characteristic of triplex DNA.[58] Of note, the CT-containing oligonucleotides targeting P1 generated an additional positive band at 295 nm, consistent with competing higher-order structure formation. To investigate the relative thermal stabilities of the stable triplexes, we performed UV melting on the complexes. For P2, the duplex melted at 64.7 ± 0.3°C, whereas triplex formation increased the melting temperature to 75.1 ± 0.1°C for the ZT-TFO and 80.1 ± 0.9°C for its LNA-modified derivative (**Figure 3c**). The *T*_m_ for the CT-containing TFO was 62.5 ± 0.5°C, which is similar to that observed for melting of the underlying duplex. For P1, the duplex melted at 75.3 ± 0.2°C, and the triplexes exhibited nearly identical *T*_m_ values of 75.8 ± 3.6°C, 74.4 ± 0.9°C, and 75.9 ± 0.4°C for the ZT-TFO, CT-TFO-clamp and ZT-TFO-clamp, respectively (**Figure 3d**). No other transitions were observed, indicating that duplex and triplex dissociated cooperatively in a single transition, as previously reported for stable triplexes.[59] The *T*_m_ for the CT-containing TFO was 59.2 ± 0.4°C at pH 5.0. Across the two panels, the order of stability was P2 ZT-TFO-LNA > P2 ZT-TFO = P1 ZT-TFO = P1 ZT-TFO-clamp > P1 and P2 CT-TFOs.

Taken together, these results demonstrate that unmodified GT-, GA-, and CT-containing TFOs are unable to form stable triplexes at neutral pH and, for the repetitive P1 sequence, are prone to competing duplex and other higher order interactions. By contrast, the ZT-containing TFOs and clamps retain high triplex specificity and robust binding under neutral pH conditions, confirming the enhanced triplex-forming capacity of the Z nucleobase relative to cytosine.

### Site-specific binding and repression of an episomal reporter

Having established stable triplex formation on synthetic duplexes, we next examined whether ZT-containing oligonucleotides could recognise their target sites within a supercoiled plasmid containing the full *MYC* promoter (pMYC). This luciferase-based construct retains the native sequence context surrounding P1 and P2 elements and has been widely used to monitor oligonucleotide-directed effects on promoter-driven expression. Since plasmid supercoiling can promote the formation of secondary DNA structures,[60] it provides a more biologically relevant substrate to assess competition between triplex formation and quadruplex formation at P1.

Triplex formation was first evaluated using a modified Cas9 protection assay in which TFO binding was expected to sterically inhibit Cas9-mediated cleavage targeted to the TFO binding sites using appropriately designed sgRNAs (**Figure S5a,b**). Cleavage efficiency was monitored by following the conversion of supercoiled (SC) plasmid to the corresponding cut linear (LIN) product.[33] Initial experiments using wild-type SpyCas9 revealed efficient cleavage at the P2 target but poor cleavage at P1 despite testing multiple sgRNAs (**Figure S5c**). The repetitive, GC-rich nature of the P1 sequence likely restricts efficient Cas9 targeting through a combination of high duplex stability, potential G-quadruplex formation, and limited PAM availability. Interestingly, switching to the PAM-relaxed SpRY Cas9 variant restored efficient cleavage at P1 using two of the four sgRNAs targeting four of the GC-tracts within the TFO target sequence on opposing strands, and this system was therefore used for subsequent protection assays (**Figure 4**).

**Figure 4.**
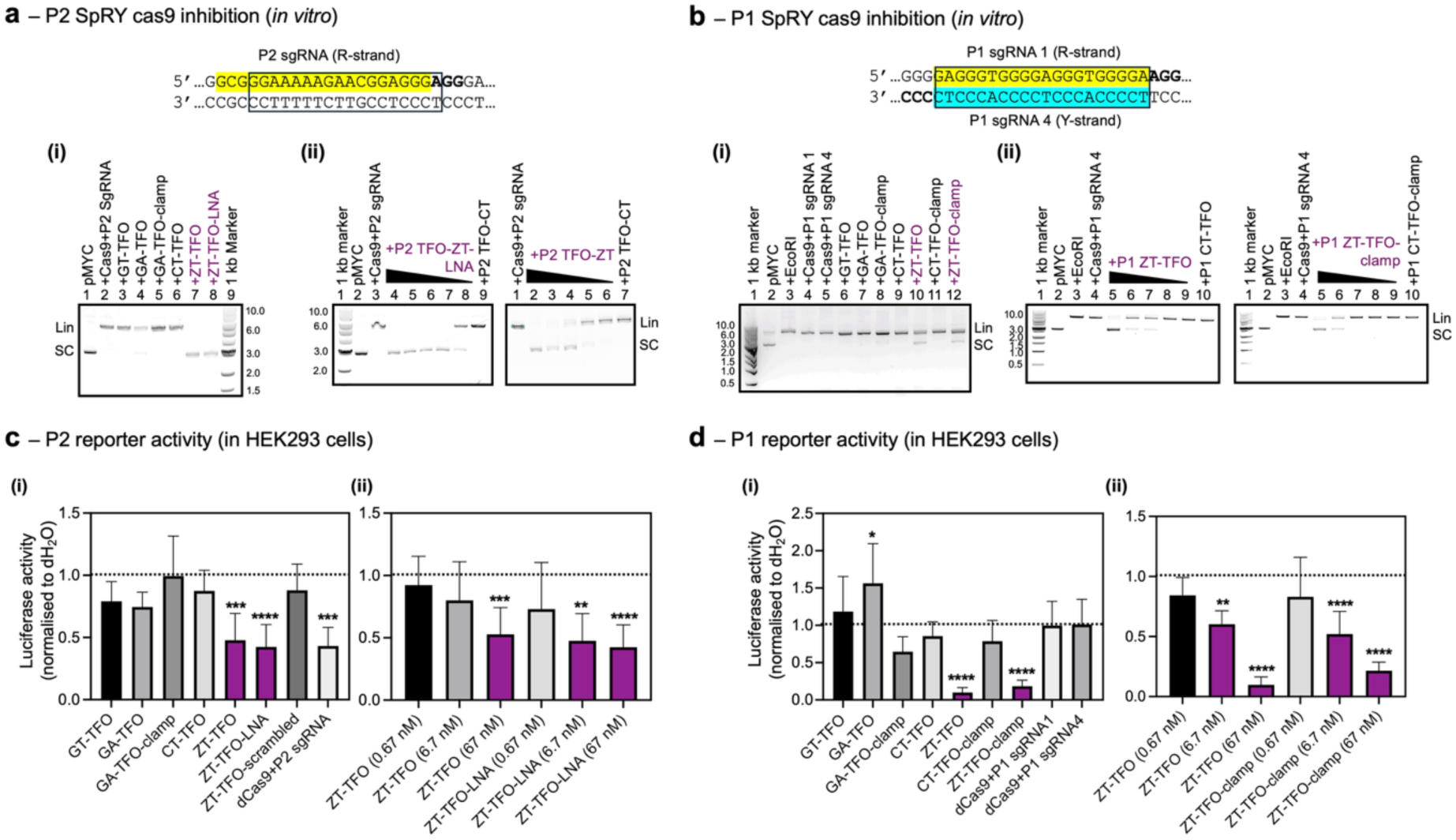
Site-specific targeting and repression at P1 and P2 in a MYC promoter-driven reporter plasmid. **(a-b)** Cas9 protection assay. sgRNA targeted either the purine (R) or pyrimidine (Y) strand (colored nucleotides) of the TFO target sequences (boxed) using PAM-relaxed SpRY Cas9. 30 nM pMYC plasmid was incubated with TFO at a final concentration of either (i) 1 µM or (ii) 10-0.1 µM in NEBuffer 3.1 buffer. The Cas9/sgRNA complex was then incubated with samples at 37°C for 1 hour. Following inactivation, cleavage products were resolved on a 0.8% agarose gel and visualized by GelRed staining. To minimize unwanted W-C interactions between P1-targeting TFOs and sgRNAs, GT- and GA-containing TFOs were evaluated using sgRNA 1, while CT- and ZT-containing TFOs were evaluated using sgRNA 4; the two sgRNAs target opposite strands of the P1 duplex. TFO binding is indicated by inhibition of Cas9-mediated cleavage of supercoiled plasmid to its linear form. **(c-d)** Effect of TFOs and catalytically inactivated dCas9 on reporter gene expression in HEK293 cells. 100 ng plasmid DNA was pre-incubated with 67 nM TFO or Cas9/sgRNA in DMEM buffer prior to co-transfection using Lipofectamine 3000^TM^. After 24 hours, luciferase activities were quantified using a SpectraMax i3x Microplate Reader. Activities were normalized to a vehicle treated dH_2_O control. Data represent at least three independent biological replicates. Statistical significance was assessed by one-way ANOVA with Tukey’s multiple-comparisons test. Error bars indicate mean ± s.d. TFO or Cas9 binding is indicated by a reduction in luciferase activity relative to control sample.

For the P2 target, the protection assay closely mirrored the EMSA data (**Figure 4a(i)**). Neither the GT-, GA-, nor CT-containing TFOs, nor the GA-TFO-clamp, inhibited SpRY Cas9 cleavage, whereas both ZT-containing TFOs afforded complete protection at 1 μM. Protection was concentration dependent and occurred at lower concentrations for the LNA-modified derivative, consistent with its greater affinity (**Figure 4a(ii)**). Because these oligonucleotides exhibited no detectable interaction with either individual target strand by EMSA, inhibition is most readily explained by triplex formation on the plasmid rather than sequestration of the sgRNA through unintended W-C interactions.

Interpretation of P1 protection required additional controls because the unmodified TFOs interacted with isolated target strands that share substantial sequence similarity with the corresponding sgRNAs. To distinguish plasmid binding from sgRNA sequestration, protection assays were performed using sgRNAs directed against either strand of the duplex. Under these conditions, only the ZT-containing TFOs inhibited Cas9 cleavage (**Figure 4b(i)**). As observed for P2, inhibition was concentration dependent (**Figure 4b(ii)**), and the relative degree of protection closely followed the order of triplex stability determined by UV melting: P2 ZT-TFO-LNA > P2 ZT-TFO > P1 ZT-TFO = P1 ZT-TFO-clamp. Together, these experiments demonstrate that ZT-containing TFOs retain sequence-selective binding within a supercoiled DNA substrate and effectively occupy their intended promoter targets.

We next determined whether triplex formation translated into repression of promoter activity following co-transfection of the pMYC plasmid and TFOs into HEK293 cells (**Figure 4c,d**). Plasmid DNA was pre-incubated with TFOs before transfection, and luciferase activity was quantified 24 h later. Based on the high affinity of the TFOs, a final oligonucleotide concentration of 67 nM was selected, substantially below that typically employed in previous cellular studies of triplex-mediated gene regulation (*ca.* 1 µM).

Targeting the P2 promoter element produced a pattern of repression that closely reflected the biophysical and protection data. Neither the GT-, GA-, nor CT-containing TFOs, nor the GA-TFO-clamp, significantly reduced reporter activity, whereas both ZT-containing TFOs decreased luciferase expression by approximately 50% (**Figure 4c(i)**). Repression was concentration dependent (**Figure 4c(ii)**) and was also not observed for Z-containing control oligonucleotide, confirming a sequence-dependent mechanism. Despite its greater affinity *in vitro*, incorporation of dispersed LNA residues did not further enhance reporter repression, suggesting that increased triplex stability alone does not necessarily increase biological activity.

The structurally more challenging P1 element produced an even greater functional response (**Figure 4d(i)**). None of the unmodified GT-, GA-, or CT-containing TFOs significantly reduced reporter activity, and the corresponding GA- and CT-containing clamp constructs were similarly inactive. In stark contrast, both the ZT-containing TFO and ZT-TFO-clamp reduced luciferase expression by approximately 80%, with repression again showing a clear concentration dependence (**Figure 4d(ii)**). As observed for P2, the higher apparent affinity of the TFO-clamp did not translate into greater biological activity. The stronger repression achieved at P1 is consistent with previous studies identifying this region as a significant regulatory element controlling *MYC* promoter activity.[54, 61]

Interestingly, the linear GA-containing TFO produced a modest, but significant, increase in reporter expression (**Figure 4d(i)**). Given its ability to bind the isolated pyrimidine strand by W-C interactions, this oligonucleotide may promote and/or stabilize strand separation, favoring G-quadruplex formation by the complementary G-rich strand,[62] although the underlying mechanism remains to be established.

For reference, catalytically inactive dCas9 ribonucleoprotein complexes were also evaluated under identical transfection conditions. At P2, dCas9 repressed reporter expression to a similar extent as the ZT-containing TFOs, whereas sgRNA alone or dCas9 in the absence of sgRNA had negligible effects (**Figure 4c(i), S6**). In contrast, neither of the two P1-targeting dCas9 complexes reduced reporter activity (**Figure 4d(i)**), consistent with the limited accessibility of this GC-rich promoter element and/or the lack of suitably positioned PAM sequences for conventional dCas9 targeting. These findings highlight the complementary advantages of triplex-mediated recognition at structurally complex promoter sequences.

Finally, potential cytotoxicity was assessed using a PrestoBlue metabolic assay (**Figure S7**). Only minor reductions in cell viability were observed across all oligonucleotide treatments, indicating that repression of reporter activity was unlikely to result from nonspecific toxicity.

Taken together, these results demonstrate that Z-containing TFOs bind site-selectively to supercoiled *MYC* promoter DNA and produce robust, sequence-dependent repression of promoter activity. Their activity closely parallels the biophysical measurements of triplex stability and extends to promoter contexts that are refractory to conventional TFOs and, in the case of P1, to dCas9-mediated regulation.

### Repression of the endogenous *MYC* promoter by ZT-containing TFOs

Having established robust triplex formation and repression of an episomal *MYC* reporter, we next investigated whether these effects translated to the endogenous *MYC* locus in HCT116 cells, which express high basal levels of MYC (**Figure S8**). Cells were transfected with TFOs using Oligofectamine under conditions optimized for oligonucleotide delivery, and endogenous *MYC* expression was quantified by qRT-PCR after 24 h and western blotting after 48 h (**Figure 5**).

**Figure 5.**
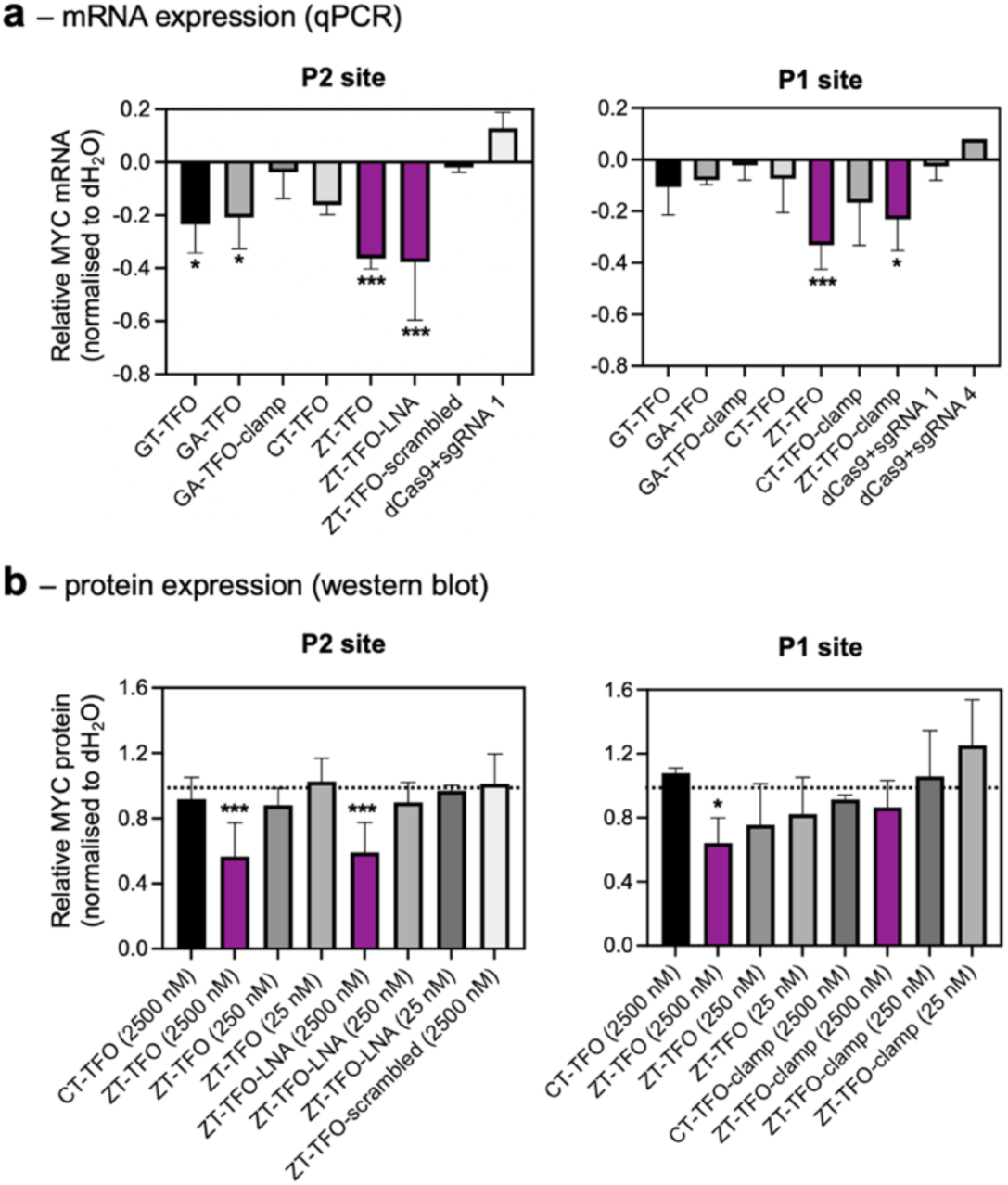
Site-specific repression at P1 and P2 in the endogenous MYC promoter in HCT116 cells. Oligonucleotides were first diluted in Opti-MEM buffer to final concentrations of 250 nM unless otherwise indicated, complexed with Oligofectamine^TM^ reagent in a total volume of 200 μl, and added to HCT116 cells. **(a)** Endogenous *MYC* RNA levels were quantified 24 h after transfection and quantified by qPCR. Total RNA was extracted, reverse transcribed, and analyzed by SYBR Green qPCR with primers specific for *MYC*, *GAPDH* and *β-actin*. Relative MYC RNA levels were normalised to *GAPDH* and *β-actin* then to the H_2_O vehicle control. **(b)** Endogenous MYC protein levels were quantified 48 h after transfection by western blotting. Cells were lysed in RIPA buffer supplemented with protease and phosphatase inhibitors, and total protein concentration was determined by BCA assay. Equal amounts of protein were resolved by SDS-PAGE, transferred to nitrocellulose membranes, and probed with antibodies against MYC and GAPDH. Signals were detected using a LI-COR Odyssey CLx imager and quantified in ImageJ. Relative MYC protein levels were normalised to GAPDH and then to the dH_2_O vehicle control. Data represent a minimum of three biological replicates; statistical significance was determined by one-way ANOVA with Tukey’s post hoc test. Triplex formation is indicated by a reduction in MYC mRNA and/or protein levels relative to control sample.

Targeting the P2 promoter element produced a pattern of repression that closely mirrored that observed in the reporter assay (**Figure 5a**). The ZT-containing TFOs produced the greatest reduction in *MYC* mRNA levels, with both the ZT-TFO and ZT-TFO-LNA decreasing transcript abundance by approximately 40-50% relative to the vehicle control. By comparison, the unmodified GT- and GA-containing TFOs produced only modest reductions, while the CT-containing TFO showed only minor activity. Importantly, the scrambled Z-containing control oligonucleotide had no significant effect on *MYC* expression, confirming that repression was sequence-dependent rather than a consequence of oligonucleotide chemistry or transfection alone.

Western blotting performed 48 h after transfection confirmed that reductions in transcript abundance were accompanied by decreases in MYC protein levels (**Figure 5b, S9**). The ZT-containing TFO consistently produced the greatest reduction in MYC protein expression, whereas the CT-containing TFO exhibited substantially weaker activity. Consistent with the qPCR data, incorporation of dispersed LNA residues did not produce a marked improvement in biological activity despite enhancing triplex stability *in vitro*, suggesting that increased intrinsic affinity alone is insufficient to increase repression at the endogenous locus.

Targeting the structurally more complex P1 promoter element produced a similar overall trend (**Figure 5a**). ZT-containing TFOs again exhibited the strongest repression among the oligonucleotide panel, whereas the corresponding unmodified GT-, GA-, and CT-containing oligonucleotides produced comparatively modest or negligible effects on endogenous *MYC* expression. Thus, despite the repetitive, G-rich nature of the P1 sequence and its propensity to adopt alternative DNA structures, incorporation of the Z nucleobase enabled efficient targeting of this challenging regulatory element in cells. Again, western blotting confirmed that reductions in transcript levels produced by the ZT-TFO and ZT-TFO-clamp were accompanied by decreases in MYC protein levels (**Figure 5b, S9**).

For comparison, catalytically inactive dCas9 ribonucleoprotein complexes directed against either the P1 or P2 promoter elements were investigated under identical conditions. In contrast to the reporter assays, no significant reduction in endogenous *MYC* expression was observed for either target site (**Figure 5a,b**). Although these experiments were optimized for oligonucleotide rather than protein delivery and therefore do not represent a direct comparison of the two technologies, they demonstrate that ZT-containing TFOs retain measurable activity under conditions where dCas9-mediated repression was not detected.

Finally, potential cytotoxicity was assessed by PrestoBlue cell viability assays performed 24 and 48 h after transfection (**Figure S10**). Only minor reductions in cell viability were observed across all oligonucleotide treatments, indicating that the reductions in *MYC* expression were unlikely to arise from nonspecific cellular toxicity.

Taken together, these results demonstrate that Z-containing parallel TFOs retain their activity at the endogenous *MYC* locus, producing robust reductions in both *MYC* mRNA and protein expression while exhibiting minimal cytotoxicity. These findings establish that the enhanced triplex-forming properties conferred by the Z nucleobase can be translated into effective regulation of endogenous gene expression in living cells.

## DISCUSSION

Sequence-specific modulation of gene expression by triplex formation has historically been limited to the antiparallel triplex-binding motif formed by purine-rich oligonucleotides.[16, 17] Although these studies established proof-of-concept for anti-gene targeting, broader application has been hindered by the relatively modest stability of this motif, often necessitating high micromolar oligonucleotide concentrations to achieve biological activity.[19, 39–46] In addition, purine-rich oligonucleotides are prone to self-association,[24, 25] formation of G-quadruplexes and related secondary structures, and sequestration of DNA-binding proteins or transcription factors through molecular decoy effects.[27] These competing interactions reduce the effective concentration of the active oligonucleotide, complicate mechanistic interpretation, and can contribute to cellular toxicity.[26] Here, we overcome these longstanding limitations by extending anti-gene targeting to the parallel triplex-binding motif using pyrimidine-rich TFOs containing the synthetic nucleobase Z.[33] While previous studies have demonstrated chromosomal binding by parallel TFOs, the oligonucleotides were highly modified and contained additional intercalating units to increase affinity and/or direct mutagenesis,[63–65] our results provide, to our knowledge, the first evidence that simple base-modified parallel-binding TFOs can directly repress expression from a native endogenous human gene.

We benchmarked our modified oligonucleotides against two well-established regulatory elements within the *MYC* promoter that differ in sequence composition, structural behavior, and biological function. The P2 element represents a more tractable target because it is less repetitive and is thought to exist predominantly as duplex DNA. Consistent with this, ZT-containing TFOs targeting P2 formed highly stable triplexes at neutral pH and reduced *MYC* promoter-driven expression by approximately 50% in both an episomal reporter assays and at the endogenous gene locus. Importantly, repression was sequence dependent, as a scrambled Z-containing control oligonucleotide showed no measurable effect on gene expression.

The P1 element, however, provided a more stringent test as this sequence contains multiple guanine tracts associated with G-quadruplex formation, and successful TFO binding is therefore likely to require competition with a dynamic duplex-quadruplex equilibrium.[49, 54] Despite this, ZT-containing TFOs formed stable triplexes with P1 and produced the strongest functional effects observed in this study, reducing reporter expression by approximately 80%, while also producing greater repression at the endogenous *MYC* locus than their unmodified counterparts. The enhanced activity observed at P1 may reflect the central role of this region in regulating MYC transcription, which has been proposed to act as a structural hub for transcriptional control.[49, 51] Triplex formation at this site may therefore inhibit transcription not only through steric occlusion, but also by altering the local structural equilibrium that governs promoter activity.

The improved biological activity of Z-containing TFOs is likely to arise from more than their enhanced triplex stability. Incorporation of the synthetic nucleobase Z reduces the propensity of oligonucleotides to self-associate or engage in unintended W-C interactions with single-stranded nucleic acids. Unlike cytosine, protonation of Z at N3 prevents stable base pairing with guanine at physiological pH while retaining Hoogsteen recognition of G-C base pairs within a triplex. This distinction was particularly evident for the repetitive P1 sequence, where conventional CT-, GT-, and GA-containing TFOs preferentially interacted with isolated target strands rather than forming stable triplexes with the intact duplex. In contrast, Z-modified TFOs showed no detectable off-target W-C interactions or higher order structure formation under the conditions examined, reducing the likelihood that transcriptional repression arose through antisense-like mechanisms, oligonucleotide self-structure, or transcription-factor sequestration. This enhanced molecular specificity is likely to be a major contributor to the improved activity observed in cells.

Comparison with dCas9 further highlights the potential utility of triplex-based targeting at GC-rich regulatory loci. Under the transfection conditions used here, dCas9 failed to repress endogenous *MYC* expression despite activity in plasmid-based assays. This likely reflects the combined challenges of delivering a large ribonucleoprotein complex to chromatinized DNA together within the nucleus and/or the reduced accessibility of repetitive, GC-rich promoter elements. These observations should not be interpreted as demonstrating that TFOs are intrinsically superior to Cas9-based approaches. Rather, they suggest that synthetic TFOs may provide complementary advantages at structurally complex genomic loci where PAM positioning, strand separation, or protein accessibility become limiting.

Interestingly, further increases in apparent triplex affinity did not translate into greater biological activity. Incorporation of LNA residues substantially increased the thermal stability of the P2 triplex, while clamp architectures enhanced binding to P1. However, neither modification produced greater repression than the corresponding ZT-containing TFOs, highlighting that nucleobase modification alone was sufficient to drive repression. One explanation is that the affinity of the unmodified ZT-TFOs already exceeds the threshold required for maximal transcriptional repression, rendering additional stabilization functionally redundant. Alternatively, improvements in intrinsic binding affinity may be offset by reduced cellular uptake, altered intracellular trafficking, or limited accessibility to chromatinized DNA, particularly for the substantially longer clamp construct. These observations suggest that, once sufficient target occupancy has been achieved, further optimization may be better directed towards improving intracellular delivery, pharmacokinetics, or expanding sequence recognition. In particular, the P1 and P2 target sites contain pyrimidine-purine inversions that could be better accommodated using nucleobase analogues that target these base pairs and may further enhance affinity and broaden the range of accessible genomic targets.[6]

An additional advantage of the parallel triplex-binding motif may be its reduced propensity to stimulate DNA repair and recombination.[23] Although the underlying mechanism remains uncertain, repair proteins are thought to recognize the helical distortions introduced by triplex formation.[66] Unlike antiparallel base triads, the Z:G-C and T:A-T triads are nearly isosteric and are therefore likely to minimize structural perturbations upon TFO binding, an effect that may be enhanced by introducing RNA-like sugar modifications that reduce the conformational rearrangements required for target binding further.[23] If so, these oligonucleotides may be particularly well suited to transient, non-mutagenic regulation of gene expression, where reversible promoter occupancy is desirable without deliberately engaging genome-modifying DNA repair pathways.

The present study focused on two regulatory elements within the *MYC* promoter, and broader design principles will require validation across a wider range of genomic targets, sequence compositions, and chromatin environments. Systematic interrogation of target-site architecture, including the contribution of individual Z:G-C base triads, inversion sites, and flanking sequence context, will help define the sequence rules governing efficient parallel triplex formation. Genome-wide occupancy and transcriptomic analyses will also be important for establishing specificity and distinguishing direct promoter repression from downstream transcriptional effects. Nevertheless, our findings establish ZT-containing parallel TFOs as a robust platform for targeting GC-rich regulatory DNA in a cellular setting. More broadly, they provide a framework for sequence-selective modulation of gene expression at genomic loci that have remained difficult to access using conventional oligonucleotides, small molecules, or protein-based targeting systems.

## DATA AVAILABILITY

Raw data has been deposited in Pure and can be accessed at https://doi.org/10.17029/a5bdcfc7-3b5d-4055-8304-fd071209c61f

## SUPPLEMENTARY DATA

Supplementary Data are available at NAR online.

## AUTHOR CONTRIBUTION

D.A.R. conceived and designed the study with input from the remaining coauthors. R.M. performed the majority of the experimental work, including biochemical, biophysical and biological characterization, and data analysis. M.B. and N.B. assisted with these experiments. H.-J.K., C.C., S.H., and S.A.B. provided modified oligonucleotides, conceptual input, and guidance. D.A.R. wrote the manuscript with input from all authors. All authors discussed the results and contributed to the final version of the manuscript. D.A.R. and S.H. supervised the project.

## FUNDING

D.R. and R.M thank the Academy of Medical Sciences (BB/L021730/1) for funding support. S.A.B and S.H thank the National Institutes of Health (R01AI135146) and the U.S. National Science Foundation (MCB-1939086) for funding support.

## CONFLICT OF INTEREST

The modified oligonucleotides described in the manuscript are commercially available from Firebird Biomolecular Sciences, LLC (www.firebirdbio.com), which operates under intellectual property owned by S.A.B. and the Foundation for Applied Molecular Evolution. Certain authors (C.C., H.J.K., and S.H.) are employed at Firebird. The remaining authors declare no competing interests.

## Notes

https://doi.org/10.17029/a5bdcfc7-3b5d-4055-8304-fd071209c61f

## REFERENCES

1. Doench JG, Hartenian E, Graham DB, Tothova Z, Hegde M, Smith I, Sullender M, Ebert BL, Xavier RJ and Root DE. Rational design of highly active sgRNAs for CRISPR-Cas9-mediated gene inactivation. Nat. Biotechnol. 2014; 32: 1262–1267.

2. Wang T, Wei JJ, Sabatini DM and Lander ES. Genetic screens in human cells using the CRISPR-Cas9 system. Science. 2014; 343: 80–84.

3. Hsu PD, Scott DA, Weinstein JA, Ran FA, Konermann S, Agarwala V, Li Y, Fine EJ, Wu X, Shalem O et al. DNA targeting specificity of RNA-guided Cas9 nucleases. Nat. Biotechnol. 2013; 31: 827–832.

4. Hoque ME, Mustafa G, Basu S and Balci H. Encounters between Cas9/dCas9 and G-quadruplexes: implications for transcription regulation and Cas9-mediated DNA cleavage. ACS Synth. Biol. 2021; 10: 972–978.

5. Balci H, Globyte V and Joo C. Targeting G-quadruplex-forming sequences with Cas9. ACS Chem. Biol. 2021; 16: 596–603.

6. Fox KR, Brown T and Rusling DA. DNA recognition by parallel triplex formation. In: Waring M (ed.). DNA-Targeting Molecules as Therapeutic Agents. Cambridge: Royal Society of Chemistry; 2018: 1–32.

7. Pozza MD, Abdullrahman A, Cardin CJ, Gasser G and Hall JP. Three’s a crowd - stabilisation, structure and applications of DNA triplexes. Chem. Sci. 2022; 13: 10193–10215.

8. Ryan K and Kool ET. Triplex-directed self-assembly of an artificial sliding clamp on duplex DNA. Chem. Biol. 1998; 5: 59–67.

9. Aviñó A, Grimau MG, Alvira M, Eritja R, Gargallo R, Orozco M and González C. Triplex formation using oligonucleotide clamps carrying 8-aminopurines. Nucleosides Nucleotides Nucleic Acids. 2007; 26: 979–983.

10. Escudé C, Garestier T and Hélène C. Padlock oligonucleotides for duplex DNA based on sequence-specific triple helix formation. Proc. Natl. Acad. Sci. U.S.A. 1999; 96: 10603–10607.

11. Prakash G and Kool ET. Structural effects in the recognition of DNA by circular oligonucleotides. J. Am. Chem. Soc. 1992; 114: 3523–3527.

12. Taladriz-Sender A, Brazzill M, Withers JM, Clark AW, Burley GA and Rusling DA. Fluorous-directed clamping stabilizes triple-helical DNA. ACS Omega. 2026; 11: 33983–33990.

13. Cecconello A, Magro M, Vianello F and Simmel FC. Rational design of hybrid DNA-RNA triplex structures as modulators of transcriptional activity in vitro. Nucleic Acids Res. 2022; 50: 13172–13182.

14. Nguyen TJD, Manuguerra I, Kumar V and Gothelf KV. Toehold-mediated strand displacement in a triplex-forming nucleic acid clamp for reversible regulation of polymerase activity and protein expression. Chem. Eur. J. 2019; 25: 12303–12307.

15. Seio K, Yamaguchi K, Yamazaki A, Kanamori T and Masaki Y. Transcription of DNA duplex containing deoxypseudouridine and deoxypseudoisocytidine, and inhibition of transcription by a triplex-forming oligonucleotide that recognizes the modified duplex. Nucleosides Nucleotides Nucleic Acids. 2020; 39: 892–904.

16. Broughton-Head VJ, Rusling DA, Fox KR and Brown T. Towards the targeted modulation of gene expression by modified triplex-forming oligonucleotides. Curr. Chem. Biol. 2008; 2: 1–10.

17. Mikame Y and Yamayoshi A. Recent advancements in development and therapeutic applications of genome-targeting triplex-forming oligonucleotides and peptide nucleic acids. Pharmaceutics. 2023; 15: 2515.

18. Psaras AM, Valiuska S, Noé V, Ciudad CJ and Brooks TA. Targeting KRAS regulation with polypurine reverse Hoogsteen oligonucleotides. Int. J. Mol. Sci. 2022; 23: 2097.

19. Valiuska S, Psaras AM, Noé V, Brooks TA and Ciudad CJ. Targeting MYC regulation with polypurine reverse Hoogsteen oligonucleotides. Int. J. Mol. Sci. 2023; 24: 378.

20. Beal PA and Dervan PB. Second structural motif for recognition of DNA by oligonucleotide-directed triple-helix formation. Science. 1991; 251: 1360–1363.

21. Durland RH, Kessler DJ, Gunnell S, Duvic M, Pettitt BM and Hogan ME. Binding of triple helix-forming oligonucleotides to sites in gene promoters. Biochemistry. 1991; 30: 9246–9255.

22. Thuong NT and Hélène C. Sequence-specific recognition and modification of double-helical DNA by oligonucleotides. Angew. Chem. Int. Ed. Engl. 1993; 32: 666–690.

23. Kalish JM, Seidman MM, Weeks DL and Glazer PM. Triplex-induced recombination and repair in the pyrimidine motif. Nucleic Acids Res. 2005; 33: 3492–3502.

24. Gellert M, Lipsett MN and Davies DR. Helix formation by guanylic acid. Proc. Natl. Acad. Sci. U.S.A. 1962; 48: 2013–2018.

25. Rippe K, Fritsch V, Westhof E and Jovin TM. Alternating d(GA) sequences form a parallel-stranded DNA homoduplex. EMBO J. 1992; 11: 3777–3786.

26. Sedoris KC, Thomas SD, Clarkson CR, Muench D, Islam A, Singh R and Miller DM. Genomic c-MYC quadruplex DNA selectively kills leukemia. Mol. Cancer Ther. 2012; 11: 66–76.

27. Cogoi S, Ballico M, Bonora GM and Xodo LE. Antiproliferative activity of a triplex-forming oligonucleotide recognizing a Ki-ras polypurine/polypyrimidine motif correlates with protein binding. Cancer Gene Ther. 2004; 11: 465–476.

28. Moser HE and Dervan PB. Sequence-specific cleavage of double-helical DNA by triple-helix formation. Science. 1987; 238: 645–650.

29. Doan TL, Perrouault L, Praseuth D, Habhoub N, Decout JL, Thuong NT, Lhomme J and Hélène C. Sequence-specific recognition, photocrosslinking and cleavage of the DNA double helix by an oligo(α-thymidylate) covalently linked to an azidoproflavine derivative. Nucleic Acids Res. 1987; 15: 7749–7760.

31. Yang Z, Hutter D, Sheng P, Sismour AM and Benner SA. Artificially expanded genetic information system: a new base pair with an alternative hydrogen-bonding pattern. Nucleic Acids Res. 2006; 34: 6095–6101.

32. von Krosigk U and Benner SA. pH-independent triple helix formation by an oligonucleotide containing a pyrazine donor-donor-acceptor base. J. Am. Chem. Soc. 1995; 117: 5361–5362.

33. Rusling DA. Triplex-forming properties and enzymatic incorporation of a base-modified nucleotide capable of duplex DNA recognition at neutral pH. Nucleic Acids Res. 2021; 49: 7256–7266.

34. Kalra S, Donnelly A, Singh N, Matthews D, Villar-Guerra RD, Bemmer V, Dominguez C, Allcock N, Cherny D, Revyakin A et al. Functionalizing DNA origami by triplex-directed site-specific photo-cross-linking. J. Am. Chem. Soc. 2024; 146: 13617–13628.

35. Brazzill M, Ma R, Munn K, Prestifilippo L, Pickford AR, Kim HJ, Chen C, Hoshika S, Benner SA and Rusling DA. Recognition of non-standard base pairs by triplex-forming oligonucleotides containing an expanded genetic alphabet. Nat. Commun. 2026; 17: 10.1038/s41467-026-74375-4

36. Gaddis SS, Wu Q, Thames HD, Digiovanni J, Walborg EF and Vasquez KM. A web-based search engine for triplex-forming oligonucleotide target sequences. Oligonucleotides. 2006; 16: 196–201.

37. Yang Z, Sismour AM, Sheng P, Puskar NL and Benner SA. Enzymatic incorporation of a third nucleobase pair. Nucleic Acids Res. 2007; 35: 4238–4249.

38. He TC, Sparks AB, Rago C, Hermeking H, Zawel L, Costa LT, Morin PJ, Vogelstein B and Kinzler KW. Identification of c-MYC as a target of the APC pathway. Science. 1998; 281: 1509–1512.

39. Kim HG and Miller DM. Inhibition of in vitro transcription by a triplex-forming oligonucleotide targeted to the human c-MYC P2 promoter. Biochemistry. 1995; 34: 8165–8171.

40. Kim HG, Reddoch JF, Mayfield C, Ebbinghaus S, Vigneswaran N, Thomas S, Jones DE and Miller DM. Inhibition of transcription of the human c-MYC proto-oncogene by intermolecular triplex formation. Biochemistry. 1998; 37: 2299–2304.

41. McGuffie EM, Pacheco D, Carbone GM and Catapano CV. Antigene and antiproliferative effects of a c-MYC-targeting phosphorothioate triplex-forming oligonucleotide in human leukemia cells. Cancer Res. 2000; 60: 3790–3799.

42. Carbone GM, McGuffie E, Napoli S, Flanagan CE, Dembech C, Negri U, Arcamone F, Capobianco ML and Catapano CV. DNA binding and antigene activity of a daunomycin-conjugated triplex-forming oligonucleotide targeting the P2 promoter of the human c-MYC gene. Nucleic Acids Res. 2004; 32: 2396–2410.

43. Catapano CV, McGuffie EM, Pacheco D and Carbone GM. Inhibition of gene expression and cell proliferation by triplex-forming oligonucleotides directed to the c-MYC gene. Biochemistry. 2000; 39: 5126–5138.

44. Boulware SB, Christensen LA, Thames HD, Coghlan L, Vasquez KM and Finch RA. Triplex-forming oligonucleotides targeting c-MYC potentiate the antitumour activity of gemcitabine in a mouse model of human cancer. Mol. Carcinog. 2014; 53: 744–752.

45. Christensen LA, Finch RA, Booker AJ and Vasquez KM. Targeting oncogenes to improve breast cancer chemotherapy. Cancer Res. 2006; 66: 4089–4094.

46. Napoli S, Negri U, Arcamone F, Capobianco ML, Carbone GM and Catapano CV. Growth inhibition and apoptosis induced by daunomycin-conjugated triplex-forming oligonucleotides targeting the c-MYC gene in prostate cancer cells. Nucleic Acids Res. 2006; 34: 734–744.

47. Cooney M, Czernuszewicz G, Postel EH, Flint SJ and Hogan ME. Site-specific oligonucleotide binding represses transcription of the human c-MYC gene in vitro. Science. 1988; 241: 456–459.

48. Levens D. How the c-MYC promoter works and why it sometimes does not. J. Natl. Cancer Inst. Monogr. 2008; 39: 41–43.

49. Siddiqui-Jain A, Grand CL, Bearss DJ and Hurley LH. Direct evidence for a G-quadruplex in a promoter region and its targeting with a small molecule to repress c-MYC transcription. Proc. Natl. Acad. Sci. U.S.A. 2002; 99: 11593–11598.

50. del Mundo IMA, Zewail-Foote M, Kerwin SM and Vasquez KM. Alternative DNA structure formation in the mutagenic human c-MYC promoter. Nucleic Acids Res. 2017; 45: 4929–4943.

51. Esain-Garcia I, Kirchner A, Melidis L, Tavares RCA, Dhir S, Simeone A, Yu Z, Madden SK, Hermann R, Tannahill D et al. G-quadruplex DNA structure is a positive regulator of MYC transcription. Proc. Natl. Acad. Sci. U.S.A. 2024; 121: e2320240121.

52. Wahlestedt C, Salmi P, Good L, Kela J, Johnsson T, Hökfelt T, Broberger C, Porreca F, Lai J, Ren K et al. Potent and nontoxic antisense oligonucleotides containing locked nucleic acids. Proc. Natl. Acad. Sci. U.S.A. 2000; 97: 5633–5638.

53. Durland RH, Rao TS, Revankar GR, Tinsley JH, Myrick MA, Seth DM, Rayford J, Singh P and Jayaraman K. Binding of T and T analogues to CG base pairs in antiparallel triplexes. Nucleic Acids Res. 1994; 22: 3233–3240.

54. Yang D and Hurley LH. Structure of the biologically relevant G-quadruplex in the c-MYC promoter. Nucleosides Nucleotides Nucleic Acids. 2006; 25: 951–968.

55. Griffin LC and Dervan PB. Recognition of thymine-adenine base pairs by guanine in a pyrimidine triple-helix motif. Science. 1989; 245: 967–971.

56. Yoon K, Hobbs CA, Koch J, Sardaro M, Kutny R and Weis AL. Elucidation of the sequence-specific third-strand recognition of four Watson-Crick base pairs in a pyrimidine triple-helix motif: T·AT, C⁺·GC, T·CG and G·TA. Proc. Natl. Acad. Sci. U.S.A. 1992; 89: 3840–3844.

57. Mergny JL, Sun JS, Rougée M, Montenay-Garestier T, Barcelo F, Chomilier J and Hélène C. Sequence specificity in triple-helix formation: experimental and theoretical studies of the effect of mismatches on triplex stability. Biochemistry. 1991; 30: 9791–9798.

58. Gray DM, Hung SH and Johnson KH. Absorption and circular dichroism spectroscopy of nucleic acid duplexes and triplexes. In: Abelson JN and Simon MI (eds). Methods in Enzymology. San Diego: Academic Press; 1995; 246: 19–34.

59. Marfurt J and Leumann CJ. Evidence for C-H···O hydrogen bond-assisted recognition of a pyrimidine base in the parallel DNA triple-helical motif. Angew. Chem. Int. Ed. 1998; 37: 175–177.

60. Belotserkovskii BP, de Silva E, Tornaletti S, Wang G, Vasquez KM and Hanawalt PC. A triplex-forming sequence from the human c-MYC promoter interferes with DNA transcription. J. Biol. Chem. 2007; 282: 32433–32441.

61. Hurley LH, Van Hoff DD, Siddiqui-Jain A and Yang D. Drug targeting of the c-MYC promoter to repress gene expression via a G-quadruplex silencer element. Semin. Oncol. 2006; 33: 498–512.

62. Rezzoug F, Thomas SD, Rouchka EC and Miller DM. Discovery of a family of genomic sequences which interact specifically with the c-MYC promoter to regulate c-MYC expression. PLoS One. 2016; 11: e0161588.

63. Brunet E, Corgnali M, Cannata F, Perrouault L and Giovannangeli C. Targeting chromosomal sites with locked nucleic acid-modified triplex-forming oligonucleotides: study of efficiency dependence on DNA nuclear environment. Nucleic Acids Res. 2006; 34: 4546–4553.

64. Faria M, Wood CD, Perrouault L, Nelson JS, Winter A, White MRH, Hélène C and Giovannangeli C. Targeted inhibition of transcription elongation in cells mediated by triplex-forming oligonucleotides. Proc. Natl. Acad. Sci. U.S.A. 2000; 97: 3862–3867.

65. Brunet E, Alberti P, Perrouault L, Babu R, Wengel J and Giovannangeli C. Exploring cellular activity of locked nucleic acid-modified triplex-forming oligonucleotides and defining its molecular basis. J. Biol. Chem. 2005; 280: 20076–20085.

66. Chin JY and Glazer PM. Repair of DNA lesions associated with triplex-forming oligonucleotides. Mol. Carcinog. 2009; 48: 389–399.

